# Bees flexibly adjust decision strategies to information content in a foraging task

**DOI:** 10.1101/2024.12.13.628313

**Authors:** Johannes Spaethe, Selma Hutzenthaler, Alexander Dietz, Karl Gehrig, James Foster, Anna Stöckl

**Affiliations:** Behavioral Physiology and Sociobiology (Zoology II), University of Würzburg, Biozentrum am Hubland, 97074 Würzburg, Germany; Department of Biology, University of Konstanz, Universitätsstr. 10, 78464 Konstanz, Germany; Zukunftskolleg, Universität Konstanz, Konstanz, Germany

**Keywords:** decision making, insect, multi-modal, foraging, trade-off, learning

## Abstract

When making decisions, humans and animals often rely on multiple sensory cues simultaneously. These provide complementary sources of information, which can help overcome ambiguity and noise, and increase the accuracy of decisions. While most studies have focused on the benefits of multimodal and within-mode integration for learning and decision making, their costs have received less attention. Processing, learning and memorizing multiple inputs requires more neural resources than for single cues, and might also require more time. In this study, we tested whether insects trade off the costs and benefits of learning multiple cues in a foraging task, using the buff-tailed bumblebee *Bombus terrestris*. To maximize comparability between cues, we presented combinations of visual-only features, as found on natural flowers: colours of varying discriminability, paired with shapes or patterns. We found that the bees relied exclusively on colours when these were easy to discriminate, and did not learn pattern or shape features presented simultaneously. With harder to discriminate colours, the bees learned both colour and shape or pattern features. Our results demonstrate that bumblebees flexibly adjust their learning strategies when presented with visual features of varying discriminability, to balance the costs and benefits of multi-cue learning. Our analysis of the learning rates of the bees with multi- and single attribute stimuli suggests that blocking could serve as a mechanism to implement this strategy switch. These results shed light on trade-offs in learning and decision making with multiple cues, and can directly be compared to studies in other insects, animals and humans.

**Significance Statement:** When making decisions, animals often rely on multiple sensory cues simultaneously. These complementary sources of information can increase decision accuracy. Unlike the benefits of cue integration, their costs have received less attention. In this study, we tested whether insects trade off the costs and benefits of learning multiple cues in a foraging task. We trained buff-tailed bumblebees, *Bombus terrestris*, to feature combinations of natural flowers. Bees flexibly switched from linear integration to winner-takes-all decision strategies depending on the sensory information available, thereby balancing accuracy and time investment in the task. We find that this switch can be explained by the cognitive phenomenon of blocking; providing the ground for future investigations into the underlying neural mechanisms of flexible decision strategies.

## Introduction

When animals and humans make decisions based on their sensory input, they often rely on multiple simultaneous cues, both within and across modalities [1–5]. We can, for example, hear and see an approaching car, and can tell a ripe from an unripe strawberry by its colour and size, scent and haptic properties. Such complementary sources of information can help overcome ambiguity when choosing between multiple options, or noise associated with one or multiple of the sensory cues [6–10], and thereby increase the accuracy of decisions in diverse species ranging from primates [11–14] to insects [15–20]. While the benefits of multimodal and within-modal integration of cues for learning and decision making are well demonstrated, the trade-offs between these benefits and potential costs have received less attention. Processing multiple inputs [21, 22], and even more so learning and memorizing them might be more costly in terms of time [23] (but see also [15, 17]), and can further lead to conflicts between the sensory inputs [24–26]. Thus, learning and integrating multiple cues might only be beneficial for decision making, when the increase in accuracy outweighs the additional costs, for example when cues are hard to detect or noisy [27–29]. When a decision can be made with high accuracy using a single cue, integrating multiple cues simultaneously might be an optimal decision-making strategy in terms of accuracy, but not in terms of efficiency [22, 30].

In this study, we tested whether insects trade off the costs and benefits of learning multiple cues in a choice task, using the buff-tailed bumblebee, *Bombus terrestris*, as a model. Insects make an ideal model system for these investigations, as they are able to take fast and accurate decisions, but possess very limited processing power [31, 32]. Much of our understanding of multimodal decision-making in insects is for visual and olfactory cues in a foraging context, for example of pollinating insects selecting and memorising flowers with visual and olfactory attributes [32–34]. Integrating cues in this context has been demonstrated to increase decision accuracy [15, 16, 18], reduce response time [15] and overcome ambiguity caused by noisy stimuli [19]. To avoid context dependent modulation as generated by olfactory cues [34, 35] and maximize comparability between cues, we presented combinations of visual-only features, as they are found on real flowers that bees visit: colours, shapes and patterns [36, 37]. We presented cue combinations which varied in the discriminability in one of the visual attributes to assess the strategies bees followed in tasks of varying difficulty.

## Results & Discussion

To test which attributes bumblebees retained from flowers displaying multiple visual cues simultaneously, we used an experimental design in which the bees first learned to associate a combination of colour and configural cue (either pattern or shape) with a sucrose reward, and a second combination of these cues with a drop of water (Fig. 1B). We then set the two attributes in conflict, by swapping them across the stimulus pair (as in [46, 47]), to test which attribute the bees relied on more strongly to make their decisions. Subsequently, bees were trained on the original cue combination again, to then test whether they had learnt the attribute they did not choose in the first test. To assess if the bees used the same learning and decision strategy in different contexts, we designed two colour categories: *distant* colour pairs and *close* colour pairs (Fig. 1C), which were easier and harder to discriminate perceptually, respectively (Fig. S1).

**Fig. 1.**
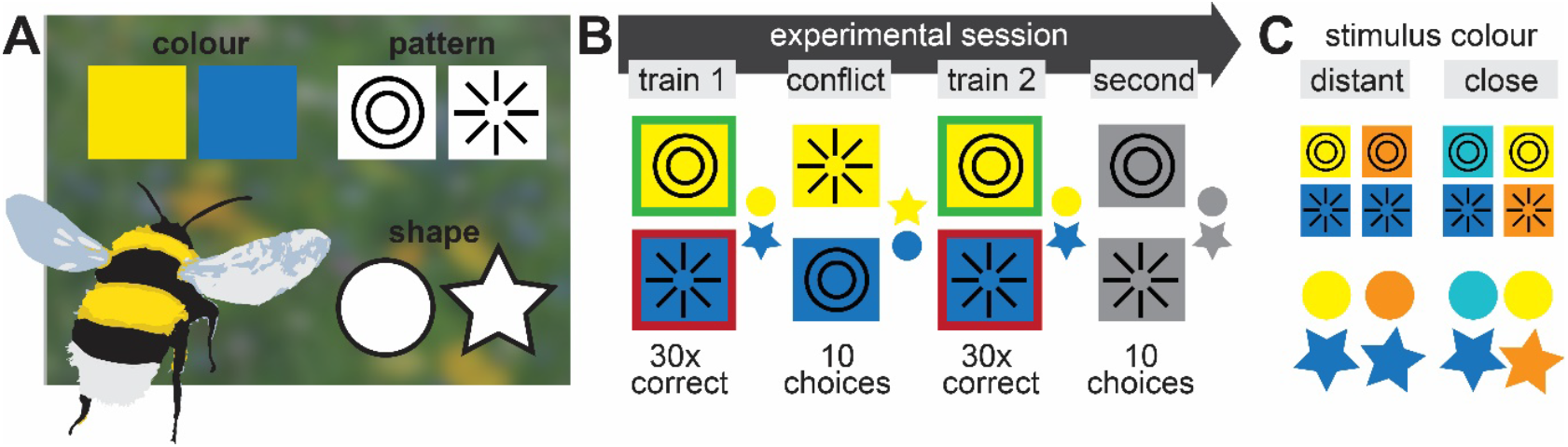
The importance of colour, pattern and shape for pollinator flower selection. **A** Flowers provide a combination of different visual cues used by insect visitors for detection and discrimination; most prominently colour, patterns and shapes. **B** To assess which of these attributes insects attend to and learn, we presented bumblebees (*Bombus terrestris*) with a combination of cues: colour and configuration (patterns or shapes). The bees were trained differentially to a colour pair, combined with either a pair of patterns or shapes, and then had to choose between the colour or configural attribute in a conflict test. Subsequently, the animals’ training was reinforced, and a second test assessed whether they learned the attribute they did not preferentially choose in the conflict test. **C** Each bee was presented with a stimulus pair out of four colour combinations for either patterns or shapes. Two of the colour pairs were distant in sensory perception, while the other two were close (see Fig. S1).

Before the training, the naïve preferences of all bees for the combination of stimuli they were trained on was assessed (Fig. S2A,B). They were then trained on the stimulus which they preferred less, to ensure that training did not reinforce innate preferences. With both configural cues, bees naively preferred blue over orange and teal over blue, irrespective of the pattern or shape the colours were combined with (Fig. S2E,F). Across colours, no significant preference for either pattern or shape emerged (Fig. S2C,D). These innate preferences also held up when stimuli were tested as single attributes (Fig. S2G).

### Bees always chose colours over other flower features when in conflict

When presented with a conflicting combination of stimuli, bees chose colour significantly more often than the shape or pattern attributes (Fig. 2A,B). The relative fraction of choices for colour depended on the colour condition: while bees chose colour over the configural attribute nearly exclusively in the *distant* colour conditions, the median choice rates for colour were only between 60–80% in the *close* colour conditions. Moreover, the individual variation was much lower with distant colours pairs, while it ranged from bees choosing pattern or shape preferentially, to those choosing colour at up to 90% with close colour pairs.

**Fig. 2.**
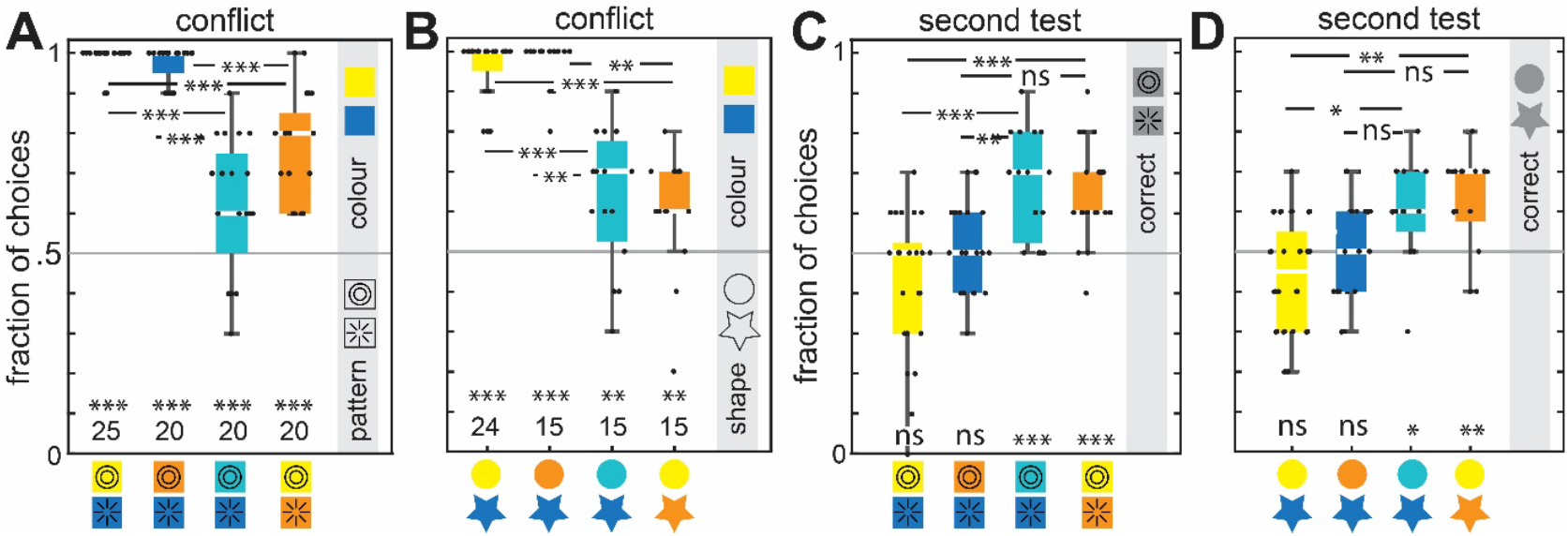
Choices for flower attributes when set into conflict. **A**,**B** Fraction of choices for the colour (above 0.5) or the pattern or shape (below 0.5) attribute in the conflict test, when the learned attributes were swapped across the stimulus pair, as compared to the training, so that each stimulus displayed only one of the two learned attributes associated with a reward. **C**,**D** Fraction of choices for the second attribute, not chosen in the conflict test (which in majority was the configural cue). For the results of animals that chose pattern or shape first, see Fig. S4B,D. Number of bees tested is indicated in A and B, for pattern and shape combinations, respectively. The results of a generalised linear mixed-effects model with a binomial family via the logit link, comparing the choice distributions to random choice at 0.5 are shown at the bottom of each graph. Comparisons across groups were performed with the same model, and are depicted where statistically significant. All statistical results are abbreviated as * < 0.05, ** < 0.01, *** < 0.001, *n*.*s*. not significant.

As in many other cases of decision making in vertebrates [25, 48] and insects [34, 49], and specifically in a flower recognition context [47, 50–53], we observed a clear hierarchy of the cues the bumblebees relied on: they primarily relied on colours over configural attributes in all conditions. Our results were highly robust, holding up qualitatively and quantitatively across multiple bumblebee colonies and different experimenters, and were quantitatively similar across the different pattern and shape features, even though these likely activate pattern and shape recognition circuits quite differently [54–58], as they varied in edge contrasts and spatial extents.

Not only was the observed hierarchy very robust, but it was also similar to that of honeybees in a comparable task with combined visual attributes. Across studies, honeybees preferentially relied on colour when recalling memorised combinations of colour, shapes or patterns [46, 59]. When cues are combined additively for decision making, the overwhelming majority of studies suggests that the more reliable, or less noisy, cues, are given more weight [3, 12, 14, 60–62]. In our experiment it seems reasonable that colours would represent the more reliable cue, since they are easier to detect at a distance than patterns and shapes [63–65], which require high resolution vision and are likely used at a more close-range stage of flower interactions [38, 66]. Moreover, under natural conditions, patterns and shapes suffer more readily from obstruction by foliage than flower colours.

### Bees learned the pattern and shape attributes only with close colour pairs

Since most of the individuals chose colour over the pattern or shape attributes in all cue combinations, we asked whether bumblebees did indeed learn both attributes of the flowers, or only the colour feature. Unlike honeybees in previous studies, which always learned the second attribute in combination training experiments [46, 59], bumblebees provided with *distant* colour pairs chose patterns or shapes at random (Fig. 2C,D), suggesting they only learned the colour, but not the pattern or shape attributes. Bees trained on the *close* colour pairs, however, chose the trained pattern or shape attribute significantly above the random choice rate (Fig. 2C,D). The few bees that did prefer the pattern or shape attribute in the conflict tests also chose the colour attributes above random choice (Fig. S2B,D), demonstrating that they learned both attributes of the combined stimuli with close colour pairs.

In previous studies with combined attributes [46, 59], honeybees learned both colour and shape or pattern attributes (though see [67] for various learning strategies with combined cues), while in our experiments, they only learned both when the two colours were perceptually close. Thus, while honeybees and bumblebees are clearly capable of learning and integrating multiple object attributes [68], and even generalise object *gestalt* across modalities [69], we here demonstrate that bumblebees do not automatically learn all object attributes, but only do so in certain contexts.

### Bees switched their decision strategy between distant and close colour pairs

We next assessed the strategy bees used to choose between cues in the conflict test for the distant and close colour conditions, in particular with respect to the weights they assigned to colour as compared with patterns or shapes, respectively. To this end, we designed a Bayesian decision model, which included the bees’ reliance on colour, pattern and shape cues, modelled as priors, which were weighted relative to each other (see Methods). In order to obtain priors for the distant and close colour conditions, as well as patterns and shapes, we trained a new set of bees to individual features, instead of cue combinations: one distant (orange–blue) and one close (teal–blue) colour pair, as well as patterns and shapes on grey background, using the same methods as for the cue combination experiment. We then tested their choices for the trained attributes (Fig. 3A). For both patterns and shapes, the resulting fractions of correct choices were not significantly different from the fraction of choices for pattern and shapes after the combined attribute training (second test, Fig. 2C,D). Qualitatively, the same holds true for the fraction of choices for close colours in the second test (Fig. S4B,D), though the small number of individuals choosing the configural category first did not allow us to confirm this statistically. This comparison demonstrates that priors derived from individual attribute training can serve as priors for the combined cue experiment.

**Fig. 3.**
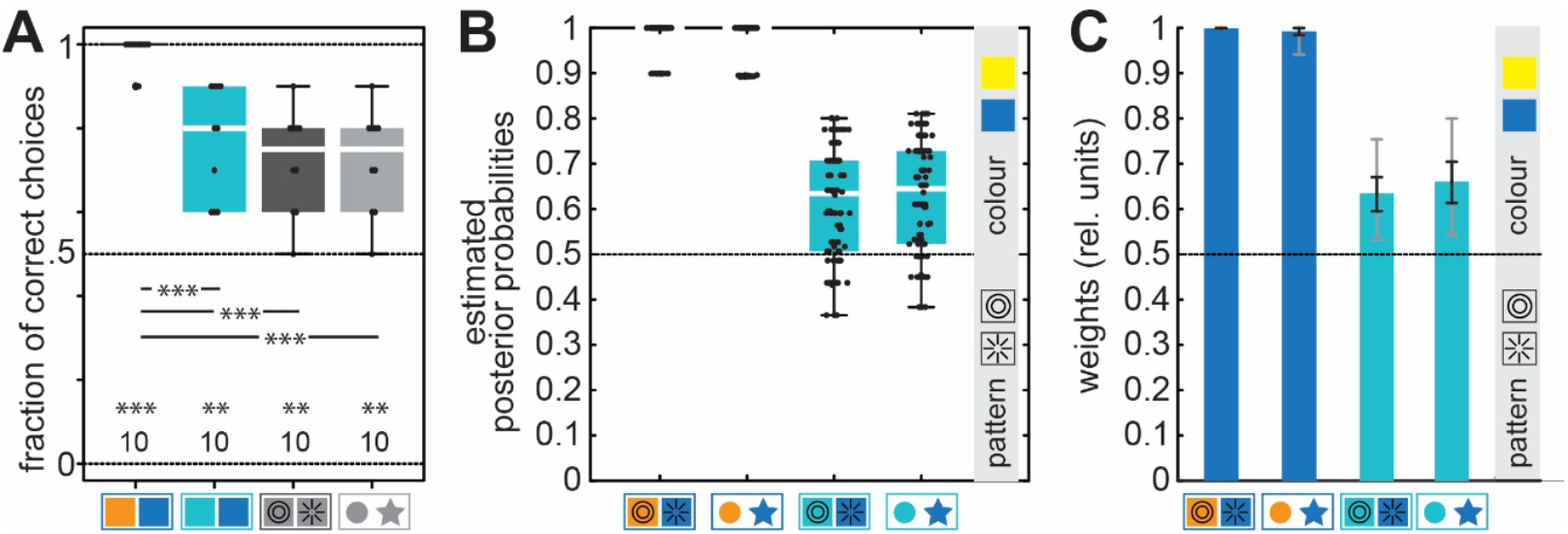
Single choices and Bayesian model with decision weights. **A** Fraction of choices after training to individual attributes (distant: orange–blue and close: blue–teal colours, patterns and shapes on grey background). The statistical results were obtained with generalised linear mixed-effects models with a binomial family using the logit link. They compare the choice fractions to random choice at 0.5, depicted at the bottom of each graph, and across conditions. All results are abbreviated as * < 0.05, ** < 0.01, *** < 0.001. **B** The predicting population choice distributions in the conflict condition, modelled as posterior probabilities using a Bayesian decision model (fitted to the corresponding experimental data shown in Fig. 2A,B), based on the experimental results for choices with individual attributes (**A**) as priors (see methods). The priors for colour and pattern or shape, respectively, were combined with a weighting factor, which was obtained by fitting the estimated posterior probabilities to the observed choices (Fig. 2A,B). **C** Estimated weighting factors, resulting from 10000 iterations of the model (depicted are mean and its 50% and 95% confidence intervals). See Fig. S3 for the distributions of weighting factors.

Using these priors, we fitted a decision model, to predict the bee’s decisions for colour in the conflict situation (Fig. 3B), in which the prior probabilities for colour and shape or pattern were linearly combined and weighted (see Methods). A weighting factor of 1 indicated that the bees only relied on their prior for colour, a factor of 0.5 that they relied on colour and patterns or shapes equally, and a factor below 0.5 that they relied more strongly on patterns or shapes than colour. The resulting weights for the four tested conditions (distant and close colours for both patterns and shapes) supported the intuitive interpretation of the conflict test results (Fig. 2A,B): the bees relied (almost) exclusively on colour with distant colour pairs (Fig. 3C), while they relied on both colour and configural cues with close colour pairs, with a stronger weighting of colour (Fig. 3C).

Combining complementary sources of information about an object is only one of two possible integration strategies – the other is a winner-takes-all approach, where the more reliable cue is selected to rely on [44, 70]. Our results suggest that the bees switch between strategies: from an additive combination of information in the close colour condition, to a winner-takes-all strategy in the distant colour condition (Fig. 3C). At an individual level, honeybees and bumblebees have been shown to trade off foraging speed for accuracy, though these were individual traits, not a switch in strategy across the population [71–75].

### Different learning rates for distant and close colour conditions suggest that blocking underlies the switch in decision strategies

In search of a mechanistic explanation for this switch in decision strategies, we analysed the learning rates of the bees with distant and close colour pairs. With combined colour and configural attributes, the bees required fewer trials to reach a high fraction of correct choices for the distant colour pairs than the close ones (Fig. 4A,B). The fraction of correct choices was already significantly higher in the first learning block and remained higher throughout all learning trials. Across combined attribute and single attribute learning of distant colours, the final learning outcome, quantified as the fraction of correct choices in the final learning block, was not significantly different (Fig. 4D).

**Fig. 4.**
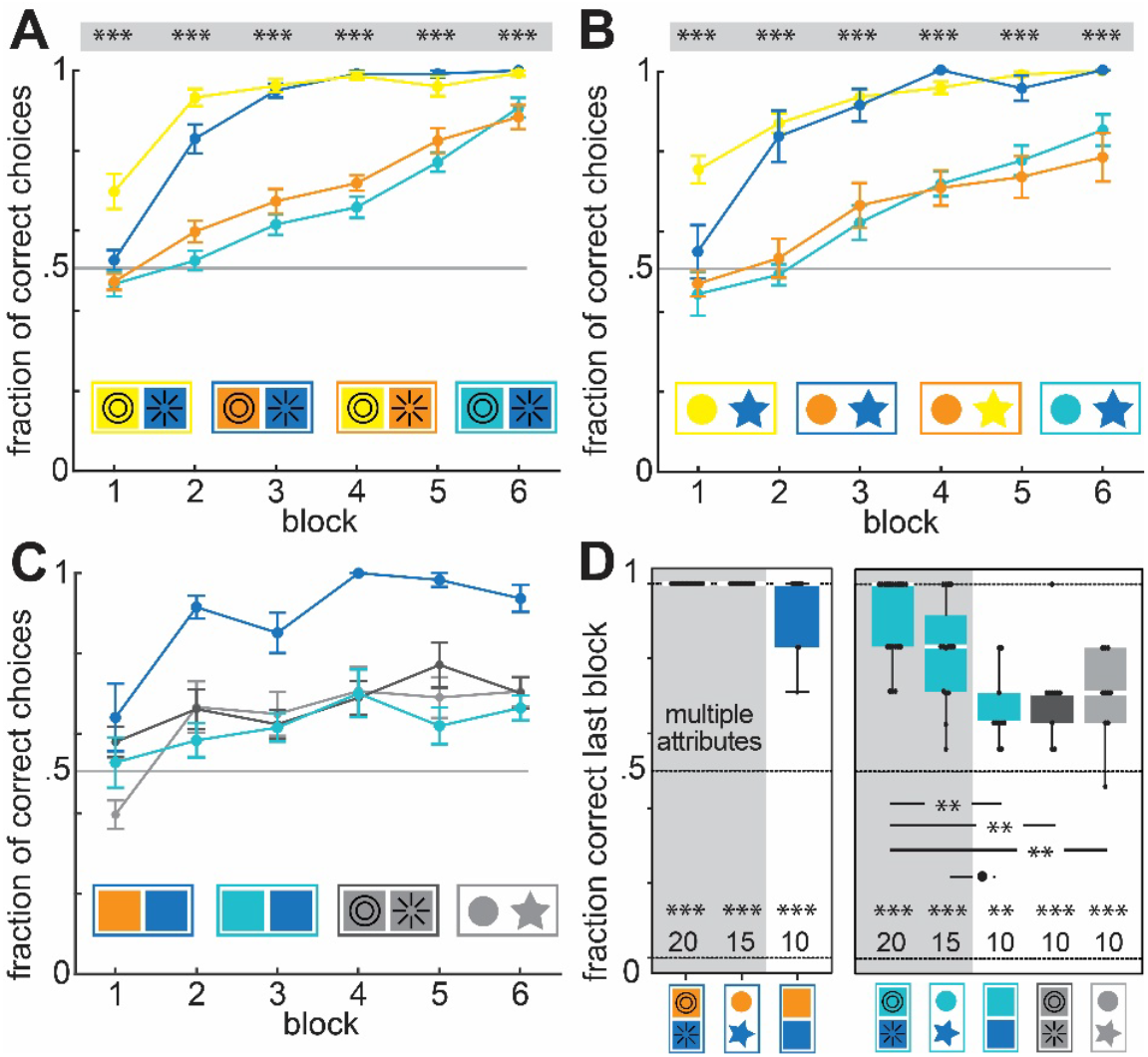
Learning performance with multi- and single attribute stimuli. **A**,**B** Learning rates of the bumblebees for all combined attributes, as well as **C** single attribute training. **D** Learning performance in the final training block, compared across multi – and single attribute learning. All statistical results were obtained with generalised linear mixed-effects models with a binomial family using the logit link, in A,B comparing the choice fractions for each block of animals with close and distant colour pairs, in D comparing the choice fractions to random choice at 0.5, depicted at the bottom of each graph. The same model was used to compare between groups, the results of which are depicted where statistically significant. All statistical results are abbreviated as * < 0.05, ** < 0.01, *** < 0.001.

The combined attribute learning rates for the close colour pairs were distinctly slower than those of the distant colour pairs (Fig. 4A,B). One might argue this was the case because the bees did learn both attributes, colour and pattern or shape (Fig. 2C,D), and therefore required more learning trials. However, the learning rates of attribute combinations with close colours were not slower than those for the respective single attributes (Fig. 4C,D). This suggests that multi-cue learning did not diminish the learning rate, but that the slower learning of the individual attributes in these conditions was responsible for the comparably slow rates. Indeed, the single attribute learning curves for close colour pairs were even flatter than their multi-attribute counterparts, and reached a significantly lower fraction of correct choices in the final learning block than the multi-attribute ones (Fig. 4D). Thus, multi-cue learning in these conditions enhanced learning rates, as well as the final learning outcome.

The difference in learning rates for distant and close colour pairs, as well as patterns and shapes, were likely linked to the discriminability of the features, and thus the complexity of the learning task. They mirrored results from previous studies which showed slower learning rates for perceptually close than for distant colours [17, 71]. The different learning rates could provide a mechanism for the change in learning and decision strategy between the two conditions: blocking [76]. Blocking is a phenomenon typically described when animals learn multiple attributes of an object or stimulus source sequentially. When one attribute is presented alone and learned, a second attribute which is presented simultaneously with the first one is not learned (as readily) any more [77, 78]. Blocking has been demonstrated across the animal kingdom, from invertebrates to vertebrates to humans. In insects, it has been described across modalities and in a variety of tasks [78–80], though the occurrence of the phenomenon can vary with the stimulus type [81] and can be confounded by experimental designs [82, 83].

In our experiments, the stimuli were not presented sequentially, as typically done when testing for blocking. But since the bees learned the distant colours much faster than shapes or patterns when presented individually (Fig. 4C), the effect might have been similar to a blocking paradigm: as the association of the reward and distant colour was already formed after a few trials, when pattern or shape associations were not yet formed, any further association of the configural attribute with the reward was blocked, as if the combination of cues had been presented only after the initial colour training. Using blocking to enact the switch in decision strategies would be a very simple, yet effective mechanism, to trade off learning speed with decision accuracy, as blocking only comes into effect when one cue is learned much faster than the other associated one. This aligns well with the general concept that the computationally limited insect brain enables simple, yet effective and adaptive strategies to perform complex behaviour, with a limited set of neurons [31, 84, 85].

### Decision strategies trade off efficiency (learning speed) and accuracy

We observed a general strategy, across all tested individuals, from an additive to a winner-takes-all approach with stimulus context, and specifically, the discriminability of one of the cues. At first glance, this might be puzzling, as one might assume that more information about a target is always better, and thus integrating complementary sources of information is optimal [7, 8]. But this does not account for time as a resource. The bees learned the distant colour pairs much faster than the close ones (Fig. 4C), which is in agreement with previous findings for learning rates with perceptually similar and distant colours [17, 71]. Interestingly, bees learned combined attributes with close colours equally fast, if not faster than when the attributes were presented individually (Fig. 4A-C). This might be due to attentional cueing [86], or valence signalling [17, 18] by one of the attributes, which are suggested to improve multicue learning rates compared to single cue ones [15, 17] (though see [23]).

Thus, with distant colour pairs, the bees could arrive at a reliable decision strategy much faster by learning only the colour attribute. Indeed, accounting for time as an important factor for decisions, instead of only the available sensory information, can change the statistically optimal decision strategy [21, 22]. This suggests that in insects as well, learning time, or efficiency, is an important factor in decision making. Further tests would be required to determine how efficiency is weighted relative to decision accuracy. This could be achieved by varying the perceptual discriminability and resulting decision accuracy of one of the cues relative to the other over a larger range, to find the point(s) at which the bees switch from a single cue to multi-cue learning and decision strategy.

### Conclusion

We found that the buff-tailed bumblebee *B. terrestris* flexibly adjusted their learning and decision strategies when presented with visual attributes of varying discriminability, to balance decision accuracy and learning efficiency. Our analysis of the bees’ learning rates suggests that blocking during learning could implement this strategy switch. This opens the door for future investigations into the underlying neural control strategies, which will enable in-depth comparisons of context-dependent learning and decision strategies in insects with the current focus in vertebrates [6, 87].

## Materials and Methods

### Animals

Colonies of *Bombus terrestris* were obtained from commercial breeders (Biobest, Westerlo, Belgium). The colonies were housed in a two-chamber wooden box (28 cm×16 cm×11 cm), which provided one chamber for nest-building and one for feeding. Before experiments started, the bees were fed ad libitum with APIinvert (Südzucker, Germany), a 70% w/v sucrose solution, and with pollen grains (Bio-Blütenpollen, Naturwaren-Niederrhein, Germany).

### Setup and Experimental Procedure

All experiments were conducted in a flight arena as previously described [38]. In brief, the bees’ nesting boxes were connected to the arena (120 cm × 100 cm × 35 cm) via an acrylic tube. The floor of the arena was covered with grey cardboard (Mi-Taintes #122, Canson SAS, Annonay Cedex, France). The same cardboard was used for the training feeders (see *Stimuli*). The experiments were performed on several different bumblebee colonies per experimental condition (see *Experimental Procedures*), to ensure that the observed effects were robust against individual colony variation [39].

### Stimuli

All artificial flowers were constructed from cardboard cut-outs, mounted on 10 mm tall, dark grey platforms 40 mm in diameter. The grey training stimuli were made from same cardboard as the floor of the arena. For testing, yellow, orange, teal and blue card were used (Tinted drawing paper #12, #19, #32 and #31, respectively; Buntpapierfabrik Ludwig Bähr, Kassel, Germany). The relative reflectance spectra for all stimuli are shown in Fig. S1A. To test the configural category *shape*, squares of 40 mm side length were used as neutral stimuli. For testing, circles of 40 mm diameter and five-armed star shapes with the same overall area were used for comparison (see Fig. 1E, insets). For the configural category *pattern*, we printed a circle-pattern consisting of two concentric black rings, and a radial pattern consisting of eight radially arranged stripes, each of 2.1 mm width onto square-shaped cardboard stimuli (see Fig. 1C, insets).

We performed two sets of experiments, in which combinations of shape-colour and pattern-colour were shown to separate groups of bumblebees. Both categories used the same pairs of colours. Two of which were *distant* in bumblebee colour space (Fig.S1), and thus should be easily distinguishable: yellow-blue, orange-blue; while two were *close*, and thus harder to distinguish: yellow-orange, teal-blue, either in combination with shapes (circle – star) or patterns (circle – radial). Neutral coloured stimuli were grey, and had a square shape in the pattern category, and star – circle shape in the shape category. Each bumblebee was only presented with one combination of colour and pattern or colour and shape, including the respective conflict and control tests (see *Experimental Procedure*).

### Spectral measurements and colour vision modelling

The relative reflectance spectra of the coloured card from which the stimuli were constructed, as well as that of the grey background on which the stimuli were presented (Fig. S1A), were measured using a JAZ spectrometer equipped with a pulsed Xenon light source (Ocean Optics, Dunedin, FL, United States). The spectrometer was calibrated against a Spectralon white diffuse reflectance standard (WS-1-SL, Ocean Optics). Colour loci in the colour triangle were calculated using the method described by [40] with using the spectral sensitivities of *B. terrestris*’ three photoreceptor classes from [41] (Fig. S1B).

### Experimental Procedure

Foragers of *B. terrestris* were selected for experiments by scoring their prior foraging activity in the flight arena on grey training flowers supplied with 30% sucrose solution. For individual identification, bees were marked on the thorax with number tags. During experiments, only a single bee was allowed to enter the arena. Before being presented with the test stimuli, five grey training stimuli with 10 µl of 50 % sucrose solution in the centre were positioned in the arena in a random arrangement. The bees performed three to five foraging bouts, separated by the forager returning to the colony, on the grey flowers. Each stimulus was refilled immediately after the bumblebee departed.

#### Conflict experiments

The experiment began with a preference test (Fig. 1B), during which the bee was presented with a combination of colour and shape or pattern (for example a yellow star and an orange circle, or a teal square with a concentric pattern and a blue square with a radial pattern). Five instances of each stimulus pair were presented in the arena, each with 10 µl of water, to avoid pairing either with a reward. The first 10 choices of each bee were recorded. Subsequently, the bee was allowed to feed ad libitum from a grey feeder and returned to the colony.

The colour–pattern or colour–shape combination of the pair that the bee least preferred was then chosen as the rewarded stimulus for training (Fig. 1B). If both stimulus pairs were preferred equally, the stimulus that had been tested on fewer individuals was selected as the rewarded one. All instances of the rewarded stimulus were supplied with 10 µl of a 1M sucrose solution, while the un-rewarded pair was supplied with the same volume of water. During training, the stimuli were re-supplied immediately after the bee approached the next stimulus. Training continued until a bee had made 30 correct choices, and as many incorrect choices as were necessary to reach 30 correct choices. When the bee returned to the colony during training, the order of the stimuli in the arena was re-arranged to prevent spatial learning. After training, sucrose solution was provided ad libitum on a grey feeder, so that the bee would fill her crop and return to the colony.

When the bee subsequently returned to the foraging arena, a conflict test was performed (Fig. 1B). It was conducted in the same way as the preference test, but the attributes of the stimuli were swapped: the rewarded colour was combined with the unrewarded pattern / shape and vice versa (for example, instead of a yellow star and an orange circle, the bee was tested with a yellow circle and an orange star).

Afterwards, a second training session was conducted using the original stimuli, until the bee again reached thirty correct choices (Fig. 1B, Fig. S4A,C).

Subsequently, a final test was conducted, which assessed the preference for the attribute the bee did not choose preferentially in the conflict test (Fig. 1B; for example, if she chose colour over shape in the conflict test, in the final test the bee was presented with grey shapes. If she chose shape over colour, the bee was presented with square-shaped coloured stimuli). This test was conducted in the same way as the preference test.

#### Single attribute experiment

This experiment was performed as above, but with stimuli that comprised only one visual attribute (distant colour: orange-blue, close colour: blue-teal, pattern or shape on grey background). The experiment began with a preference test comprised of ten unrewarded choices, followed by a training session until thirty correct choices were performed, and an unrewarded session, in which the stimulus pair, which was presented for training, was shown unaltered.

### Number of bees and colonies per experiment

For the colour and pattern conflict experiment, bumblebee foragers of four colonies were trained and tested by two different experimenters (25 individuals in the yellow–blue, and 20 each in the orange–blue, teal–blue, and yellow–orange conditions), while four colonies were tested by three different experimenters for the colour and shape experiment (24 individuals in the yellow–blue, and 15 each in the orange–blue, teal–blue, and yellow–orange conditions). Two additional colonies were used to test 10 foragers in each condition with single attribute stimuli by one experimenter.

### Bayesian decision model

We constructed a Bayesian decision model [42, 43], predicting the choice rates of bees for colour in the conflict test as posterior probabilities p(H∣O), which are generated by the likelihood of the observations given the hypothesis p(O∣H), and the prior probability of the hypothesis p(H), which entails the animals’ prior knowledge about the probability of a reward on a given stimulus.

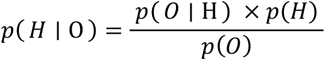

The likelihood of the observations given the hypothesis p(O∣H) was 0.5 in all experiments, since we presented 2 choices of stimuli in equal numbers. The term p(O) provides a normalising term, to retain the resulting probability within the bounds of [0,1], calculated as:

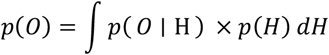

Since the decision model was designed to predict choices in the conflict situation, the priors needed to reflect both prior probability of colour p(H∣C_1_) and pattern or shape p(H∣C_2_), respectively. We therefore generated a combined prior, with a relative weighting (weighting factor w) between the two terms [12, 44, 45]:

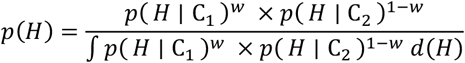

The combined prior also included a normalising term, which integrates all possible prior probabilities for H. The full model thus reads:

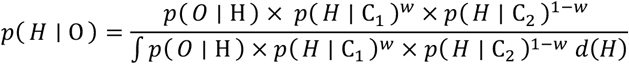

We implemented the model in discrete terms in Matlab 2022a (The Mathworks, Natick, US), and fitted the resulting posterior probabilities to the observed choices of the animals in the conflict test, to obtain the weighting factor for each of the four conditions. As priors, we assigned the fraction of choices for the rewarded colour or pattern/ shape obtained in the individual attribute experiments, in which the animals were only trained two a single attribute (distant or close colour, pattern, shape), instead of a colour combination (Fig. 3A). Since the single attribute data was obtained from a different set of bumblebees than those that performed the conflict experiments, we used a permutation approach: every choice data point for the colour attribute was combined with every point for the pattern or shape attribute to calculate the combined prior, resulting in a 10×10 prior matrix (as 10 bees were tested in both single attribute colour and pattern or shape tests). From the resulting 100 posterior probabilities, we randomly selected a subset matched to the number of observed choices in the respective conflict conditions (15 or 20). For these, a weighting factor was fitted, using a sum of mean square error between the posterior probabilities and observed choice frequencies (using the fmincon function with bounds at 0 and 1 and a maximum of 10^6^ iterations). This procedure was repeated 10000 times for different subsets, with each model run initiated with a randomised starting parameter drawn from a Gaussian distribution centred on 0.5, to arrive at the same number of weighting factor estimates for each of the four conditions (Fig. 3C, Fig. S4). Posterior probabilities were calculated for the mean of the weighting factor distribution, using the full 10×10 posterior matrix (Fig. 3B), for visual inspection of the goodness of fit.

### Data analysis

To assess learning progress, the fraction of correct choices was calculated for each block of five correct choices, and as many incorrect choices as were made to arrive at the default value (Fig. 1C,D). This assured equal possibility for forming a positive association to the stimulus across bees. For all unrewarded preference tests, the fraction of correct choices was calculated. Statistical analysis was performed using with R v4.1.2 (R Foundation for Statistical Computing, Vienna, Austria).

To assess whether the preference for colour in the conflict experiment differed from random choice (50:50, since all stimuli were presented in pairs), a generalised linear mixed-effects model with the following formula was used:

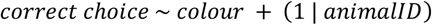

These estimated the fixed-effect of the stimulus colour on the relative probability of choosing the stimulus – scored for each individual as 0 or 1, accounting for individual biases.

To analyse whether the preference for pattern or shape, if they were not preferred in the conflict experiment, differed from random choice (50:50, since all stimuli were presented in pairs) in the final test, a generalised linear mixed-effects model with the following formula was used:

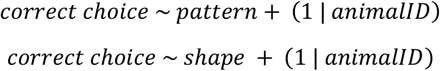

These estimated the fixed-effect of the stimulus pattern or shape on the relative probability of choosing the stimulus – scored for each individual as 0 or 1, accounting for individual biases.

To compare the choice fraction for colour in the conflict test, and the non-preferred attribute in the final test for patterns and shapes, we used a generalised linear model of the family “quasibinomial”:

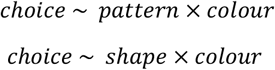

To assess the fraction of choices of naïve bees for colour, shape or pattern, and the effects of their interaction, we used a generalised linear model of the family “quasibinomial”:

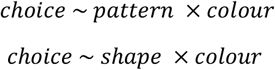

## Author Contributions

Conceptualization: J.S., A.S.; Methodology: J.S., A.S., J.F.; Validation: J.S., A.S., J.F.; Formal analysis: J.S., A.S.; Investigation: S.H., A.D., K.G.; Resources: J.S., A.S.; Data curation: A.S., S.H., A.D., K.G.; Writing - original draft: A.S.; Writing - review & editing: J.S., A.S., J.F.; Visualization: A.S.; Supervision: J.S., A.S.; Funding acquisition: A.S.

## Acknowledgements

We acknowledge funding to A.S. by the Bavarian Academy of Sciences and Humanities, the Zukunftskolleg Konstanz, and the Emmy Noether Programme of the DFG (STO 1255/4-1).

## Data Availability

Source data for all behavioural experiments and spectral measurements: https://figshare.com/s/c4c912de7dd7e24a9ec8

Custom-written analysis code: https://github.com/stoeckl-lab/Spaethe_et_al_2024_beeDecisions

## Literature

[1] Raposo, D., Sheppard, J.P., Schrater, P.R., and Churchland, A.K. (2012). Multisensory Decision-Making in Rats and Humans. The Journal of Neuroscience 32, 3726–3735.

[2] Guilford, T. and Dawkins, M.S. (1991). Receiver psychology and the evolution of animal signals. Animal Behaviour 42, 1–14.

[3] Angelaki, D.E., Gu, Y., and DeAngelis, G.C. (2009). Multisensory integration: psychophysics, neurophysiology, and computation. Current Opinion in Neurobiology 19, 452–458.

[4] Hebets, E.A. and Papaj, D.R. (2004). Complex signal function: developing a framework of testable hypotheses. Behavioral Ecology and Sociobiology 57, 197–214.

[5] Rubi, T.L. and Stephens, D.W. (2016). Does multimodality per se improve receiver performance? An explicit comparison of multimodal versus unimodal complex signals in a learned signal following task. Behavioral Ecology and Sociobiology 70, 409–416.

[6] Coen, P., Sit, T.P., Wells, M.J., Carandini, M., and Harris, K.D. (2023). Mouse frontal cortex mediates additive multisensory decisions. Neuron 111, 2432–2447.e13.

[7] Seilheimer, R.L., Rosenberg, A., and Angelaki, D.E. (2014). Models and processes of multisensory cue combination. Current Opinion in Neurobiology 25, 38–46.

[8] Jones, P.R. (2016). A tutorial on cue combination and Signal Detection Theory: Using changes in sensitivity to evaluate how observers integrate sensory information. Journal of Mathematical Psychology 73, 117–139.

[9] Partan, S.R. (2017). Multimodal shifts in noise: switching channels to communicate through rapid environmental change. Animal Behaviour 124, 325–337.

[10] Wilson, A.J., Dean, M., and Higham, J.P. (2013). A game theoretic approach to multimodal communication. Behavioral Ecology and Sociobiology 67, 1399–1415.

[11] Sumby, W.H. and Pollack, I. (1954). Visual Contribution to Speech Intelligibility in Noise. The Journal of the Acoustical Society of America 26, 212–215.

[12] Ernst, M.O. and Banks, M.S. (2002). Humans integrate visual and haptic information in a statistically optimal fashion. Nature 415, 429–433.

[13] Kuang, S. and Zhang, T. (2014). Smelling directions: Olfaction modulates ambiguous visual motion perception. Scientific Reports 4.

[14] Fetsch, C.R., Turner, A.H., DeAngelis, G.C., and Angelaki, D.E. (2009). Dynamic Reweighting of Visual and Vestibular Cues during Self-Motion Perception. The Journal of Neuroscience 29, 15601–15612.

[15] Kulahci, I.G., Dornhaus, A., and Papaj, D.R. (2008). Multimodal signals enhance decision making in foraging bumble-bees. Proceedings of the Royal Society B: Biological Sciences 275, 797–802.

[16] Leonard, A.S. and Masek, P. (2014). Multisensory integration of colors and scents: insights from bees and flowers. Journal of Comparative Physiology A 200, 463–474.

[17] Gil-Guevara, O., Bernal, H.A., and Riveros, A.J. (2022). Honey bees respond to multimodal stimuli following the principle of inverse effectiveness. Journal of Experimental Biology 225.

[18] Okray, Z., Jacob, P.F., Stern, C., Desmond, K., Otto, N., Talbot, C.B., Vargas-Gutierrez, P., and Waddell, S. (2023). Multisensory learning binds neurons into a cross-modal memory engram. Nature 617, 777–784.

[19] Jordan, K.A., Sprayberry, J.D., Joiner, W.M., and Combes, S.A. (2024). Multimodal processing of noisy cues in bumblebees. iScience 27, 108587.

[20] Buehlmann, C., Mangan, M., and Graham, P. (2020). Multimodal interactions in insect navigation. Animal Cognition 23, 1129–1141.

[21] Crevecoeur, F., Munoz, D.P., and Scott, S.H. (2016). Dynamic Multisensory Integration: Somatosensory Speed Trumps Visual Accuracy during Feedback Control. The Journal of Neuroscience 36, 8598–8611.

[22] Drugowitsch, J., DeAngelis, G.C., Klier, E.M., Angelaki, D.E., and Pouget, A. (2014). Optimal multisensory decision-making in a reaction-time task. eLife 3, e03005.

[23] Strube-Bloss, M., Günzel, P., Nebauer, C.A., and Spaethe, J. (2023). Visual accelerated and olfactory decelerated responses during multimodal learning in honeybees. Frontiers in Physiology 14.

[24] Parisi, G.I., Barros, P., Kerzel, M., Wu, H., Yang, G., Li, Z., Liu, X., and Wermter, S. (2017). A computational model of crossmodal processing for conflict resolution. In 2017 Joint IEEE International Conference on Development and Learning and Epigenetic Robotics (ICDL-EpiRob) (IEEE), pp. 33–38.

[25] De Gelder, B. and Bertelson, P. (2003). Multisensory integration, perception and ecological validity. Trends in Cognitive Sciences 7, 460–467.

[26] Song, Y.H., Kim, J.H., Jeong, H.W., Choi, I., Jeong, D., Kim, K., and Lee, S.H. (2017). A Neural Circuit for Auditory Dominance over Visual Perception. Neuron 93, 940–954.e6.

[27] Pecher, D. and Zeelenberg, R. (2022). Does multisensory study benefit memory for pictures and sounds? Cognition 226, 105181.

[28] Munoz, N.E. and Blumstein, D.T. (2019). Optimal multisensory integration. Behavioral Ecology 31, 184–193.

[29] Rhebergen, F., Taylor, R.C., Ryan, M.J., Page, R.A., and Halfwerk, W. (2015). Multimodal cues improve prey localization under complex environmental conditions. Proceedings of the Royal Society B: Biological Sciences 282, 20151403.

[30] Fawcett, T.W. and Johnstone, R.A. (2003). Optimal assessment of multiple cues. Proceedings of the Royal Society of London. Series B: Biological Sciences 270, 1637–1643.

[31] Chittka, L. and Niven, J. (2009). Are bigger brains better? Curr. Biol. 19, R995–R1008.

[32] Barron, A.B., Gurney, K.N., Meah, L.F.S., Vasilaki, E., and Marshall, J.A.R. (2015). Decision-making and action selection in insects: inspiration from vertebrate-based theories. Frontiers in Behavioral Neuroscience 9.

[33] Giurfa, M., Núñez, J., and Backhaus, W. (1994). Odour and colour information in the foraging choice behaviour of the honeybee. Journal of Comparative Physiology A 175.

[34] Thiagarajan, D. and Sachse, S. (2022). Multimodal Information Processing and Associative Learning in the Insect Brain. Insects 13, 332.

[35] Zhang, L.Z., Zhang, S.W., Wang, Z.L., Yan, W.Y., and Zeng, Z.J. (2014). Cross-modal interaction between visual and olfactory learning in Apis cerana. Journal of Comparative Physiology A 200, 899–909.

[36] Leonard, A.S., Dornhaus, A., and Papaj, D.R. (2011). Forget-me-not: Complex floral displays, inter-signal interactions, and pollinator cognition. Current Zoology 57, 215–224.

[37] Orbán, L.L. and Plowright, C.M. (2014). Getting to the start line: how bumblebees and honeybees are visually guided towards their first floral contact. Insectes Soc. 61, 325–336.

[38] Richter, R., Dietz, A., Foster, J., Spaethe, J., and Stöckl, A. (2023). Flower patterns improve foraging efficiency in bumblebees by guiding approach flight and landing. Funct Ecol 37, 763–777.

[39] Maharaj, G., Horack, P., Yoder, M., and Dunlap, A.S. (2018). Influence of preexisting preference for color on sampling and tracking behavior in bumble bees. Behavioral Ecology 30, 150–158.

[40] Chittka, L. and Kevan, P. (2005). Practical Pollination Biology, chapter Flower colors as advertisement. (Cambridge, Canada.: Enviroquest), pp. 157–206.

[41] Briscoe, A.D. and Chittka, L. (2001). The evolution of color vision in insects. Annual Review of Entomology 46, 471–510.

[42] Berger, J.O. (1985). Statistical Decision Theory and Bayesian Analysis (Springer New York).

[43] Jaynes, E.T. (2003). Probability Theory: The Logic of Science (Cambridge University Press).

[44] Clark, J.J. and Yuille, A.L. (1990). Data Fusion for Sensory Information Processing Systems (Springer US).

[45] Yuille, A. and Bülthoff, H. (1996). Bayesian decision theory and psychophysics (Cambridge University Press), pp. 123–162.

[46] Gould, J.L. and Towne, W.F. (1988). Honey Bee Learning (Elsevier), pp. 55–86.

[47] Stöckl, A., Heinze, S., Charalabidis, A., El Jundi, B., Warrant, E., and Kelber, A. (2016). Differential investment in visual and olfactory brain areas reflects behavioural choices in hawk moths. Sci. Rep. 6, 26041.

[48] Meijer, G.T., Mertens, P.E., Pennartz, C.M., Olcese, U., and Lansink, C.S. (2019). The circuit architecture of cortical multisensory processing: Distinct functions jointly operating within a common anatomical network. Progress in Neurobiology 174, 1–15.

[49] Wessnitzer, J. and Webb, B. (2006). Multimodal sensory integration in insects - towards insect brain control architectures. Bioinspiration & Biomimetics 1, 63–75.

[50] Kriston, I. (1973). Die Bewertung von Duft-und Farbsignalen als Orientierungshilfen an der Futterquelle durchApis mellifera L. Journal of Comparative Physiology 84, 77–94.

[51] Kinoshita, M., Stewart, F.J., and Ômura, H. (2017). Multisensory integration in Lepidoptera: Insights into flower-visitor interactions. BioEssays 39.

[52] Balkenius, A. and Kelber, A. (2006). Colour preferences influences odour learning in the hawkmoth, Macroglossum stellatarum. Naturwissenschaften 93, 255–258.

[53] Dacke, M., Bell, A.T.A., Foster, J.J., Baird, E.J., Strube-Bloss, M.F., Byrne, M.J., and el Jundi, B. (2019). Multimodal cue integration in the dung beetle compass. Proceedings of the National Academy of Sciences 116, 14248–14253.

[54] Srinivasan, M. (1994). Pattern recognition in the honeybee: Recent progress. Journal of Insect Physiology 40, 183–194.

[55] Heisenberg, M. (1995). Pattern recognition in insects. Curr. Opin. Neurobiol. 5, 475–481.

[56] Dill, M. and Heisenberg, M. (1995). Visual pattern memory without shape recognition. Phil. Trans. R. Soc. Lond. B 349, 143–152.

[57] Horridge, A. (2009). Generalization in visual recognition by the honeybee (Apis mellifera). Journal of Insect Physiology 55, 499–511.

[58] Chen, L., Zhang, S., and Srinivasan, M.V. (2003). Global perception in small brains: Topological pattern recognition in honey bees. Proc. Natl. Acad. Sci. U.S.A. 100, 6884–6889.

[59] Hsu, P.S. and Yang, E.C. (2012). The critical cue in pattern discrimination for the honey bee: Color or form? Journal of Insect Physiology 58, 934–940.

[60] Fetsch, C.R., Pouget, A., DeAngelis, G.C., and Angelaki, D.E. (2011). Neural correlates of reliability-based cue weighting during multisensory integration. Nature Neuroscience 15, 146–154.

[61] Körding, K.P. and Wolpert, D.M. (2004). Bayesian integration in sensorimotor learning. Nature 427, 244–247.

[62] Chandrasekaran, C. (2017). Computational principles and models of multisensory integration. Current Opinion in Neurobiology 43, 25–34.

[63] Menzel, R., Gumbert, A., Kunze, J., Shmida, A., and Vorobyev, M. (1997). Pollinator’s strategies in finding flowers. Israel Journal of Plant Sciences 45, 141–156.

[64] de Ibarra, N.H., Langridge, K.V., and Vorobyev, M. (2015). More than colour attraction: behavioural functions of flower patterns. Curr. Opin. Insect Sci. 12, 64–70.

[65] Hempel de Ibarra, N., Holtze, S., Bäucker, C., Sprau, P., and Vorobyev, M. (2022). The role of colour patterns for the recognition of flowers by bees. Philosophical Transactions of the Royal Society B: Biological Sciences 377.

[66] Kannegieser, S., Kraft, N., Haan, A., and Stöckl, A. (2024). Visual guidance fine-tunes probing movements of an insect appendage. Proceedings of the National Academy of Sciences 121, e2306937121.

[67] Horridge, A. (2007). The preferences of the honeybee (Apis mellifera) for different visual cues during the learning process. Journal of Insect Physiology 53, 877–889.

[68] Rands, S.A., Whitney, H.M., and Hempel de Ibarra, N. (2023). Multimodal floral recognition by bumblebees. Current Opinion in Insect Science 59, 101086.

[69] Solvi, C., Gutierrez Al-Khudhairy, S., and Chittka, L. (2020). Bumble bees display cross-modal object recognition between visual and tactile senses. Science 367, 910–912.

[70] Battaglia, P.W., Jacobs, R.A., and Aslin, R.N. (2003). Bayesian integration of visual and auditory signals for spatial localization. Journal of the Optical Society of America A 20, 1391.

[71] Dyer, A. and Chittka, L. (2004). Bumblebees (Bombus terrestris) sacrifice foraging speed to solve difficult colour discrimination tasks. Journal of Comparative Physiology A 190.

[72] Chittka, L., Dyer, A.G., Bock, F., and Dornhaus, A. (2003). Bees trade off foraging speed for accuracy. Nature 424, 388–388.

[73] Wang, M.Y., Chittka, L., and Ings, T.C. (2018). Bumblebees Express Consistent, but Flexible, Speed-Accuracy Tactics Under Different Levels of Predation Threat. Frontiers in Psychology 9.

[74] Ings, T.C. and Chittka, L. (2008). Speed-Accuracy Tradeoffs and False Alarms in Bee Responses to Cryptic Predators. Current Biology 18, 1520–1524.

[75] Burns, J.G. and Dyer, A.G. (2008). Diversity of speed-accuracy strategies benefits social insects. Current Biology 18, R953–R954.

[76] Kamin, L.J. (1969). Punishment and Aversive Behavior, chapter Predictability Surprise, Attention, and Conditioning (New York: Appleton-Century-Crofts), pp. 279–296.

[77] Prados, J., Alvarez, B., Acebes, F., Loy, I., Sansa, J., and Moreno-Fernández, M.M. (2013). Blocking in rats, humans and snails using a within-subjects design. Behavioural Processes 100, 23–31.

[78] Terao, K., Matsumoto, Y., and Mizunami, M. (2015). Critical evidence for the prediction error theory in associative learning. Scientific Reports 5.

[79] Couvillon, P.A., Arakaki, L., and bitterman, M.E. (1997). Intramodal blocking in honeybees. Animal Learning &amp; Behavior 25, 277–282.

[80] Hosler, J.S. and Smith, B.H. (2000). Blocking and the detection of odor components in blends. Journal of Experimental Biology 203, 2797–2806.

[81] Guerrieri, F., Lachnit, H., Gerber, B., and Giurfa, M. (2005). Olfactory blocking and odorant similarity in the honeybee. Learning &amp; Memory 12, 86–95.

[82] Blaser, R.E., Couvillon, P.A., and Bitterman, M.E. (2006). Blocking and pseudoblocking: New control experiments with honeybees. Quarterly Journal of Experimental Psychology 59, 68–76.

[83] Blaser, R.E., Couvillon, P.A., and Bitterman, M.E. (2008). Within-subjects experiments on blocking and facilitation in honeybees (Apis mellifera). Journal of Comparative Psychology 122, 373–378.

[84] Guiraud, M.G., Gallo, V., Quinsal-Keel, E., and MaBouDi, H. (2025). Bumblebee visual learning: simple solutions for complex stimuli. Animal Behaviour, 221: 123070

[85] MaBouDi, H., Barron, A.B., Li, S., Honkanen, M., Loukola, O.J., Peng, F., Li, W., Marshall, J.A.R., Cope, A., Vasilaki, E., et al. (2021). Non-numerical strategies used by bees to solve numerical cognition tasks. Proceedings of the Royal Society B: Biological Sciences 288, 20202711.

[86] Eckstein, M.P., Mack, S.C., Liston, D.B., Bogush, L., Menzel, R., and Krauzlis, R.J. (2013). Rethinking human visual attention: Spatial cueing effects and optimality of decisions by honeybees, monkeys and humans. Vision Research 85, 5–19.

[87] Pagan, M., Tang, V.D., Aoi, M.C., Pillow, J.W., Mante, V., Sussillo, D., and Brody, C.D. (2024). Individual variability of neural computations underlying flexible decisions. Nature 10.1038/s41586-024-08433-6.

