## Supplement for "Bees flexibly adjust decision strategies to information content in a foraging task"

### Supplementary Figures

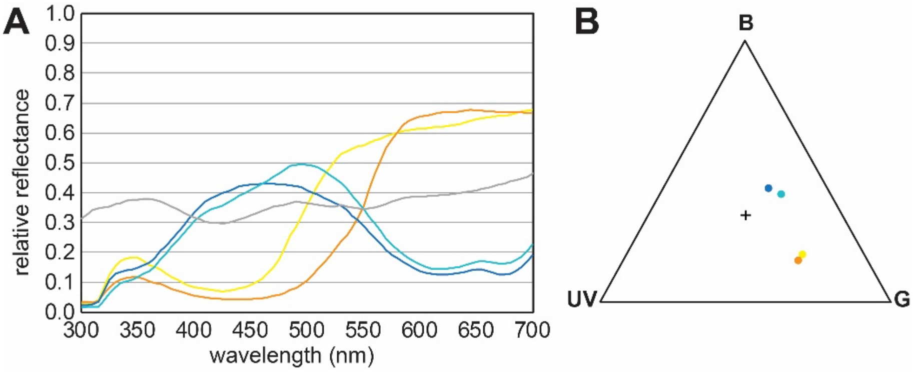


**Fig. S1 Stimulus colour spectra and relative photoreceptor activation.**
**A** Relative reflectance spectra of the coloured carton used to construct the four colours of the stimuli, as well as the grey carton used as a background and to construct the neutral pattern and shape stimuli. **B** Loci of the four tested colours plotted in a colour triangle assuming receptor adaptation of the three photoreceptor types (UV=ultraviolet,B=blue, G=green) of bumblebees to the grey background (see Methods).

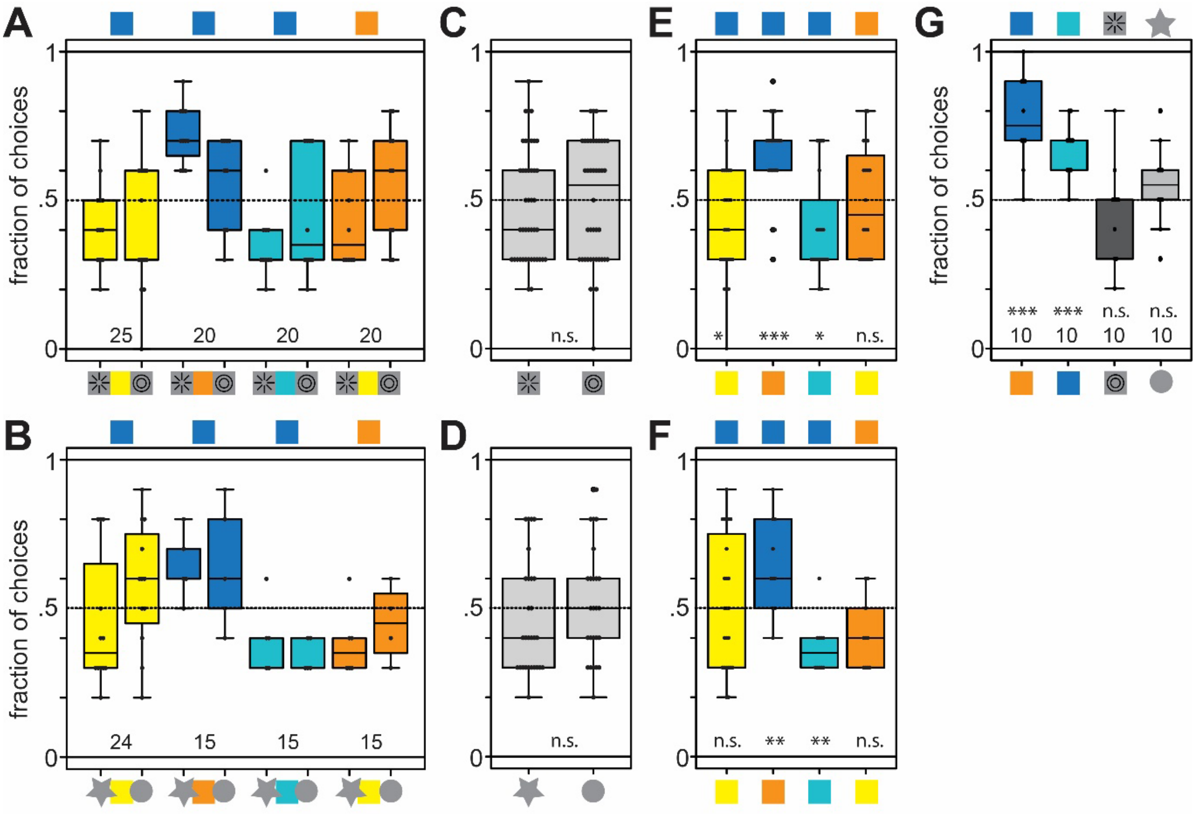


**Fig. S2 Naïve preferences of the bumblebees for colours, patterns and shapes.**
**A,B** Naïve preferences of the bees in all experiments with cue combinations, measured in 10 unrewarded choices, with one stimulus pair each per bee (number of bees for all combinations at the bottom of each panel). **C,D** Preferences for radial and concentric patterns, and star and circle shapes, respectively, across all colour pairs. **E** Preferences within each of the four colour combinations, across patterns and shapes, respectively. The statistical results in A-E were obtained with generalised linear mixed-effects models with a binomial logit link. The full model with interaction between patterns and colour, or shape and colour, did not have a significantly lower deviance than the model with only colour. E compare the choice fractions to random choice at 0.5, depicted at the bottom of each graph. **G** Naïve preferences of the bees before the single attribute learning trials. **H** Fraction of correct choices in an

unrewarded test after single attribute learning. The statistical results in G,H were obtained with generalised linear mixed-effects models with a binomial logit link. They compare the choice fractions to random choice at 0.5, depicted at the bottom of each graph, and across conditions for H. All statistical results are abbreviated as * < 0.05, ** < 0.01, *** < 0.001, n.s. not significant.

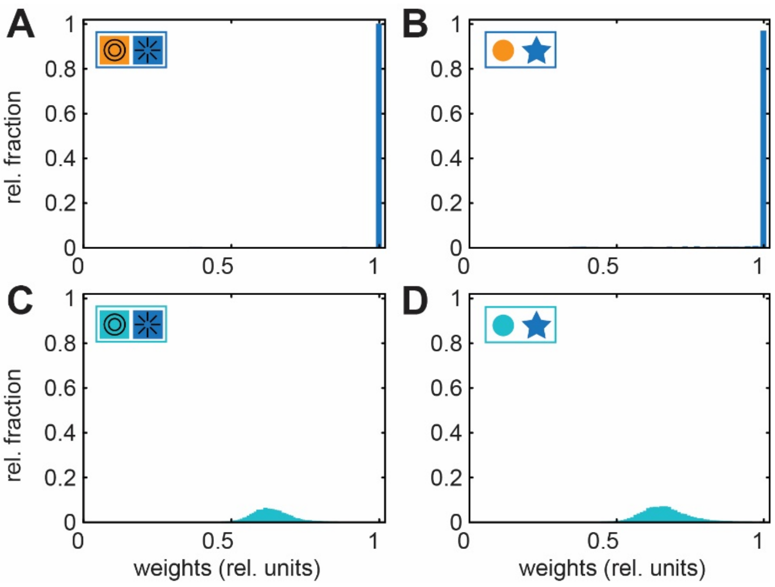


**Fig. S3 Distribution of decision weights resulting from Bayesian decision model.**
**A-D** Distribution of weighting factors, which were obtained by fitting a Bayesian decision model to the conflict data of the orange and blue, and teal and blue conditions (see Methods, Fig. 3). The distributions resulted from 10000 iterations of the model.

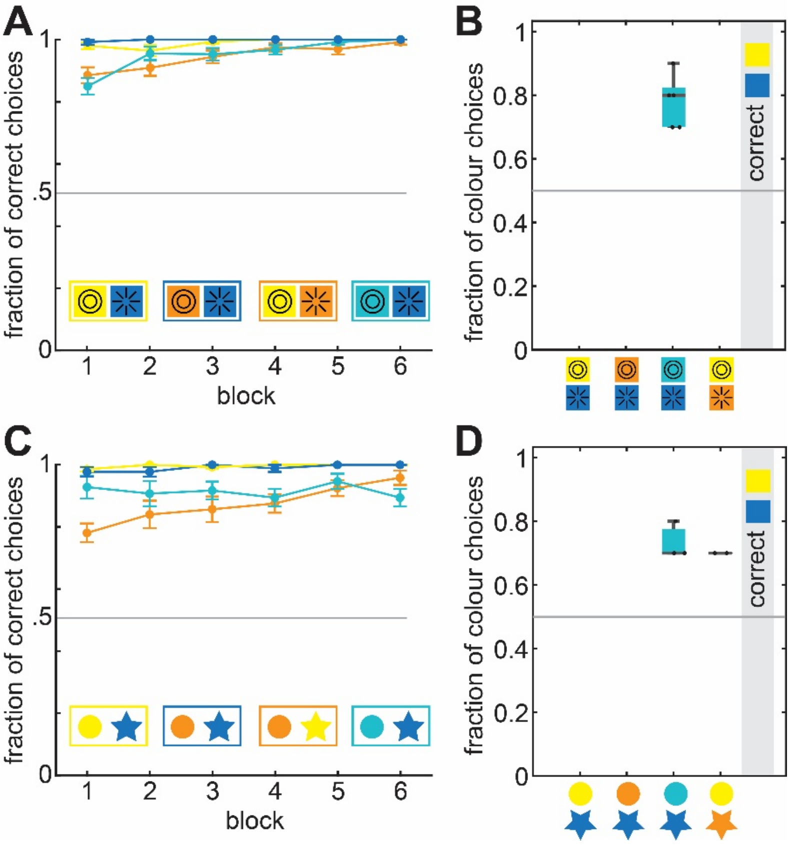


**Fig. S4 Learning rates of the second training, and colour results of the second attribute test.**

**A,C** Learning rates of bumblebees in the second training session for all pattern and shape conditions.
**C,D** Fraction of correct choices of bees that chose patterns, or shapes, as their first attribute in the conflict test.
